# Chorioamnionitis Is a Risk Factor for Intraventricular Hemorrhage in Preterm Infants: A Systematic Review and Meta-Analysis

**DOI:** 10.1101/334375

**Authors:** Eduardo Villamor-Martinez, Monica Fumagalli, Owais Mohammed Rahim, Sofia Passera, Giacomo Cavallaro, Pieter Degraeuwe, Fabio Mosca, Eduardo Villamor

## Abstract

Although chorioamnionitis (CA) is a well-known risk factor for white matter disease of prematurity, the association with intraventricular hemorrhage (IVH) is controversial and has not been yet systematically reviewed. We performed a systematic review and meta-analysis of studies exploring the association between CA and IVH. A comprehensive literature search was conducted using PubMed/MEDLINE and EMBASE, from their inception to 1 July 2017. Studies were included if they examined preterm infants and reported primary data that could be used to measure the association between exposure to CA and the presence of IVH. A random-effects model was used to calculate odds ratios (OR) and 95% confidence intervals (CI). We found 1284 potentially relevant studies, of which 85 met the inclusion criteria (46,244 infants, 13,432 CA cases). Meta-analysis showed that CA exposure was significantly associated with all grades IVH (OR 1.88, 95% CI 1.61-2.19), with grades 1-2 IVH (OR 1.69, 95% CI 1.22-2.34), and with grades 3-4 IVH (OR 1.62, 95% CI 1.42-1.85). Both clinical and histological CA were associated with an increased risk for developing IVH in very preterm infants. In contrast, the presence of funisitis did not increase IVH risk when compared to CA in the absence of funisitis (OR 1.22, 95% CI 0.89-1.67). Further meta-analyses confirmed earlier findings that CA-exposed infants have significantly lower gestation age (GA; mean difference [MD] −1.20 weeks) and lower birth weight (BW; MD −55g) than the infants not exposed to CA. However, meta-regression and subgroup analysis could not demonstrate an association between the lower GA and BW and the risk of IVH in the CA-exposed infants. In conclusion, our data show that CA is a risk factor for IVH, but also a risk factor for greater prematurity and more clinical instability. In contrast to other complications of prematurity, such as patent ductus arteriosus, retinopathy of prematurity, or bronchopulmonary dysplasia, the effect of CA on IVH appears to be independent of CA as causative factor for very preterm birth.

## 1. Introduction

Germinal matrix hemorrhage–intraventricular hemorrhage (GMH-IVH) is one of the most common complications of prematurity (1–3). IVH typically initiates in the germinal matrix, which is a richly vascularized collection of neuronal-glial precursor cells in the developing brain and may disrupt the ependymal lining and extend into the lateral ventricle (1–3). Severe IVH (grade 3-4) is associated with increased mortality as well as short- and long-term neurological morbidity, whilst the short-term and long-term outcomes of milder forms of IVH (grade 1-2) are less established, and they remain a significant research area (1, 2, 4).

As extensively reviewed by Inder et al., the pathogenesis of IVH is multifactorial and may involve intravascular, vascular, and extravascular factors (2). Intravascular factors relate to the regulation of blood flow, pressure, and volume in the microvascular bed of the germinal matrix as well as to platelet-capillary function and blood clotting capability (2). Vascular factors refer to the intrinsic fragility and vulnerability of germinal matrix blood vessels (2). Extravascular factors include the poor support of the extravascular space surrounding the germinal matrix capillaries, the postnatal decrease in extravascular tissue pressure, and an excessive fibrinolytic activity (2). As assessed by Inder et al., not all the pathogenetic factors are present in every IVH and the clinical circumstances determine which factors are most relevant in each infant (2). Among these clinical circumstances, very preterm birth, generally defined as birth before 32 completed weeks of gestation, is the most consistently associated with the development of IVH. However, a number of risk factors including, among others, absent antenatal corticosteroid (ACS) treatment, vaginal delivery, peri- and postnatal hypoxic-ischemic events, severe respiratory distress syndrome (RDS), pneumothorax, hypercapnia, hemodynamic disturbances (either systemic hypertension or hypotension), rapid volume expansion, decreased hematocrit, glucose and/or electrolyte disturbances, seizures, patent ductus arteriosus (PDA), thrombocytopenia, inherited thrombophilia, and infection may predispose to the development of IVH (1–3, 5–8).

Several studies suggest that IVH is unequally distributed among the different leading causes of very preterm delivery (9, 10). An estimated 40% of very preterm births are associated with placental inflammation, which is often subclinical. This inflammation may be localized to the maternal placenta or membrane (chorioamnionitis) or may extend to the fetus, inducing an inflammatory response, which is evidenced by funisitis (11–16). Chorioamnionitis (CA) is not only a major risk factor for (very) preterm birth, but it is also considered a major risk factor for the morbidity and mortality associated with prematurity (11–16). The pathogenetic role of CA in the development of complications of prematurity, such as necrotizing enterocolitis (NEC), bronchopulmonary dysplasia (BPD), PDA, retinopathy of prematurity (ROP), or cerebral palsy has been addressed in several systematic reviews (17–23). Although intrauterine inflammation is a well-known risk factor for white matter disease of prematurity (24), the association with IVH is controversial and has not been yet systematically reviewed. Moreover, a consideration with any analysis of CA as a risk factor for preterm morbidity, is accounting for the role of GA, birth weight (BW) and other baseline characteristics which differ between CA-exposed and CA-unexposed infants (17, 20, 22, 23). With this in mind, we aimed to perform a systematic review and meta-analysis of studies exploring the association between CA and IVH, as well as the role of potential confounding factors.

## 2. Methods

The methodology followed the same structure as earlier meta-analyses on CA and ROP (22), and CA and PDA (20). We developed a protocol a priori, which specified the objectives, inclusion criteria, method for evaluating study quality, included outcomes and covariates, and statistical methodology. We report the study according to the guidelines for Preferred Reporting Items for Systematic Reviews and Meta-Analysis (PRISMA) (25).

### 2.1 Sources and search strategy

We performed a comprehensive literature search in the PubMed/MEDLINE and EMBASE databases from their inception to July 1, 2017. The search strategy involved combining the following keywords in various ways: “chorioamnionitis”, “intrauterine infection” “intrauterine inflammation”, “antenatal infection” “antenatal inflammation” “intraventricular hemorrhage”, “risk factors”, “outcome”, “cohort”, and “case-control”. No studies were excluded based on language. In addition, we used the following strategies to identify additional studies: review of reference lists of previous systematic reviews on CA, and of articles included in the present review, and the use of the “cited by” tool in Web of Science and Google Scholar.

### 2.2 Study selection

We included studies which evaluated infants who were preterm (<37 weeks) or low BW (<2500g), as well as studies which used stricter inclusion criteria. Studies were included if they reported primary data on the association between CA-exposure and IVH. We included studies which reported the rate of IVH in infants with and without CA, and studies which reported the rate of CA in infants with and without IVH. The results of the total search were screened independently by two reviewers (O.M.R, E.V.), in several rounds: first by title only, second by title and abstract, and thirdly by consulting the full text. The reviewers resolved discrepancies in inclusion through discussion and by consulting a third reviewer (P.D).

### 2.3 Data extraction

Using a predetermined worksheet, two researchers (E.V.-M., O.M.R.) extracted data from the studies included. Another two investigators (P.D., E.V.) checked the extracted data for accuracy and completeness. We resolved discrepancies by discussion and through checking the primary report. The following data were extracted from each study: citation information, location of study, primary objective, criteria for inclusion/exclusion of infants, definitions used for CA and for IVH, infant baseline characteristics in the total group and the CA-exposed and CA-unexposed groups, and reported results on the outcomes of interest (including raw numbers, summary statistics and adjusted analyses on CA and IVH where available).

### 2.4 Quality assessment

We used the Newcastle-Ottawa Scale (NOS) for cohort or case-control studies to assess the methodological quality of included studies. Three aspects of a study are evaluated by the NOS: selection, comparability and exposure/outcome, and these are scored individually and tallied up to a total of 9 points. Two researchers (E.V.-M. and E.V.) independently used the NOS to evaluate the quality of each study, and discrepancies were discussed and resolved by consensus.

### 2.5 Statistical Analysis

We combined and analyzed studies using comprehensive meta-analysis V 3.0 software (CMA, RRID:SCR_012779, Biostat Inc., Englewood, NJ, USA). We calculated the odds ratio (OR) and 95% confidence intervals (CI) for dichotomous outcomes from the data extracted from the studies. We calculated the mean difference (MD) and 95% CI for continuous outcomes. We used the method of Wan and colleagues (26) to estimate the mean and standard deviation, when continuous variables were reported as median and range/interquartile range in studies. We used a random-effects model to calculate summary statistics, due to anticipated heterogeneity. This method accounts for both intra-study and inter-study variability.

A mixed-effects model was used for subgroup analyses (27). This model is characterized by a random-effects model that combines studies within subgroups, and a fixed-effects model that combines subgroups together to create an overall effect. This model does not assume that study-to-study variance (tau-squared) is the same in all subgroups. We assessed statistical heterogeneity using the Cochran’s Q statistic, which reflects the degree of variance, and the I^2^-statistic, which describes the proportion of observed variance that is due to variance in true effect sizes rather than sampling error (28). Visual inspection of funnel plots and Egger’s regression test were used to evaluate evidence of publication bias.

We used univariate random-effects meta-regression (method of moments) to evaluate covariates which may affect the effect size (29). We defined the following covariates a priori as potential sources of variability: CA type (clinical or histological), funisitis, differences in GA and BW between the infants with and without CA, use of ACS, mode of delivery, rate of preeclampsia, rate of small for gestational age (SGA), rate of premature rupture of membranes (PROM), rate of RDS, rate of PDA, rate of early onset sepsis (EOS), rate of late onset sepsis (LOS) and mortality. A probability value of less than 0.05 (0.10 for heterogeneity) was considered statistically significant. We considered probability values under 0.05 (0.10 for heterogeneity) as statistically significant.

## 3. Results

### 3.1 Description of studies

After removing duplicates, we found 1284 potentially relevant studies, of which 85 (7, 30–113) met the inclusion criteria. Figure 1 depicts the PRISMA flow diagram of the search. The included studies evaluated 46,244 infants, including 13,432 cases of CA. The characteristics of the included studies are summarized in Supplementary Table 1. Fifty-eight studies examined the outcomes of maternal CA and included IVH as one of the outcomes. Twenty-four studies evaluated risk factors for developing IVH and included maternal CA as of the risk factors. Five studies studied the association between CA and IVH as their primary outcome (35, 72, 93, 113, 114). Fifty-four studies used a histological definition of CA and 24 studies used a clinical definition of CA. Only two studies (55, 61) examined microbiological CA and IVH. One study (77) provided data on IVH and its association with histological and clinical CA separately. In four studies (48, 53, 62, 109) infants were assigned to the CA group if they presented histological and/or clinical CA.

**Figure 1.**
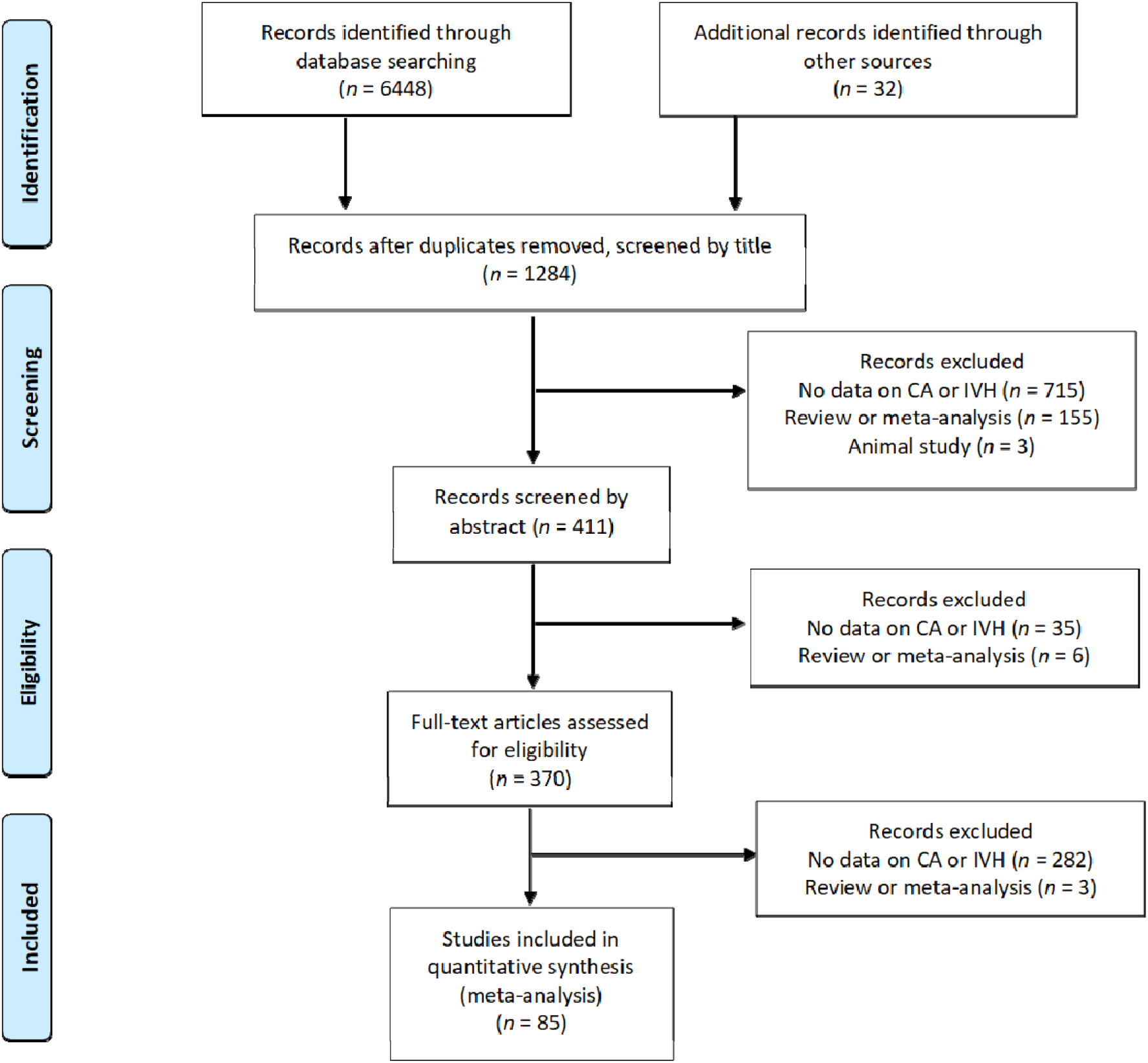
Flow diagram of search process.

### 3.2 Quality assessment

A summary of the quality assessment of each study using the NOS is shown in Supplementary Table 2. One study received a quality score of 5 points, 19 studies achieved a quality score of six points, 43 studies achieved a quality score of seven points, 11 studies achieved a quality score of eight, and 11 studies achieved the maximum score of 9 points. Studies were downgraded in quality for not adjusting the risk of IVH for confounders (*k* = 62), for not defining IVH clearly (*k* = 9), for only adjusting the risk of IVH for one confounding factor (*k* = 7), for not defining CA clearly (*k* = 6), and for losing a substantial portion of infants to follow-up (*k* = 4).

### 3.3 Analysis based on unadjusted data

Meta-analysis showed that CA exposure was significantly associated with all grades IVH (Figure 2A), with grades 2-4 IVH (Figure 2B), with grades 1-2 IVH (2C), and with grades 3-4 IVH (Figure 2D). When the type of CA was analyzed separately, histological CA remained significantly associated with all grades IVH (Figure 3), with grades 2-4 IVH (Figure 2B), with grades 1-2 IVH (Figure 2C), and with grades 3-4 IVH (Figure 4). Clinical CA was significantly associated with all grades IVH (Figure 5) and with grades 3-4 IVH (Figure 6), but not with grades 1-2 IVH (Figure 2C). There was only one study providing data on the association of clinical CA and IVH grades 2-4 (Figure 2B). We could not find significant evidence of publication bias through visual inspection of the funnel plot (Figure 7), or through Egger’s regression test.

**Figure 2.**
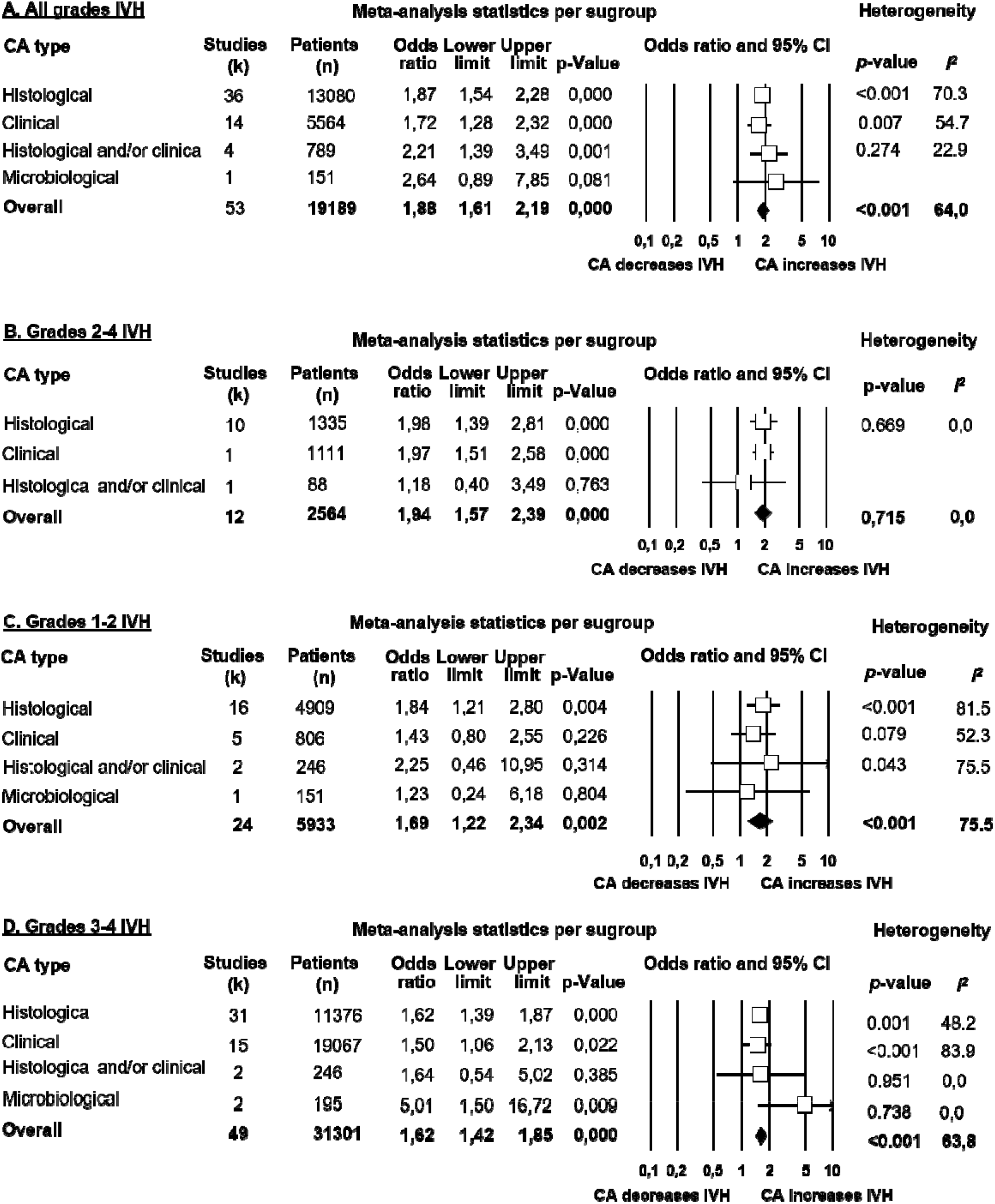
Meta-analyses of the association between chorioamnionitis (CA) and intraventricular hemorrhage (IVH), according to definition of IVH. CI: confidence interval.

**Figure 3.**
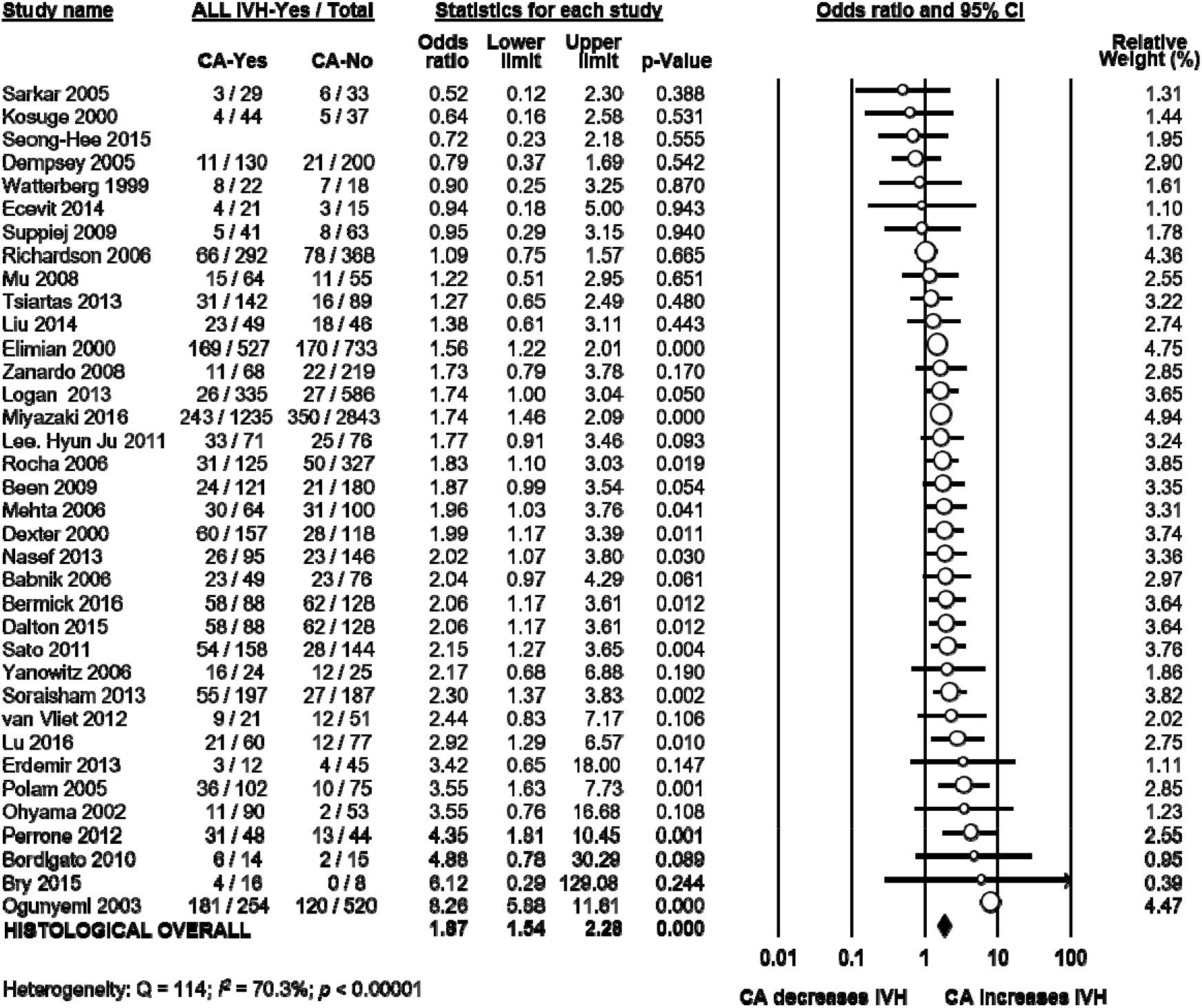
Meta-analysis of the association between histological chorioamnionitis (CA) and all grades intraventricular hemorrhage (IVH). CI: confidence interval.

**Figure 4.**
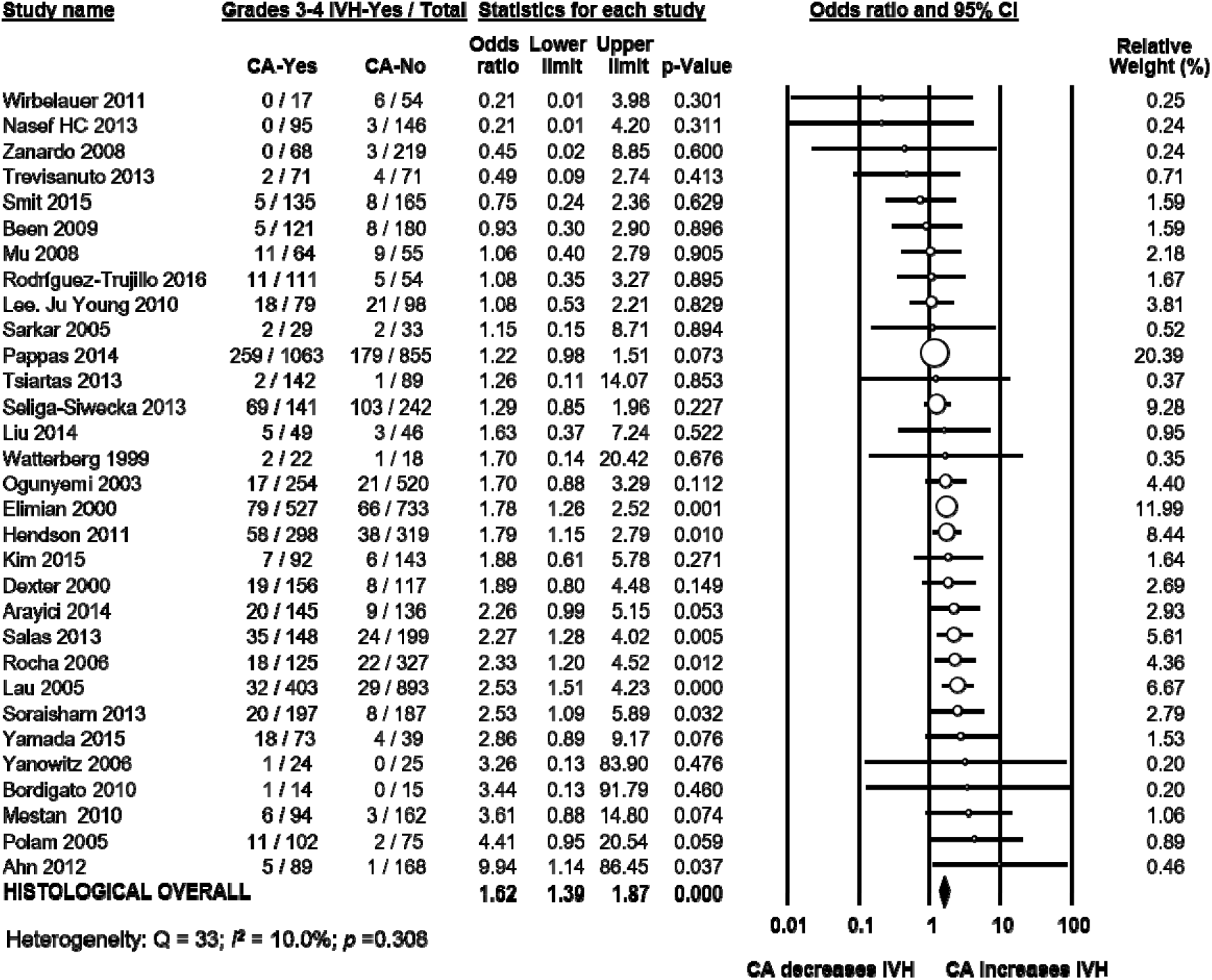
Meta-analysis of the association between histological chorioamnionitis (CA) and grades 3-4 intraventricular hemorrhage (IVH).

**Figure 5.**
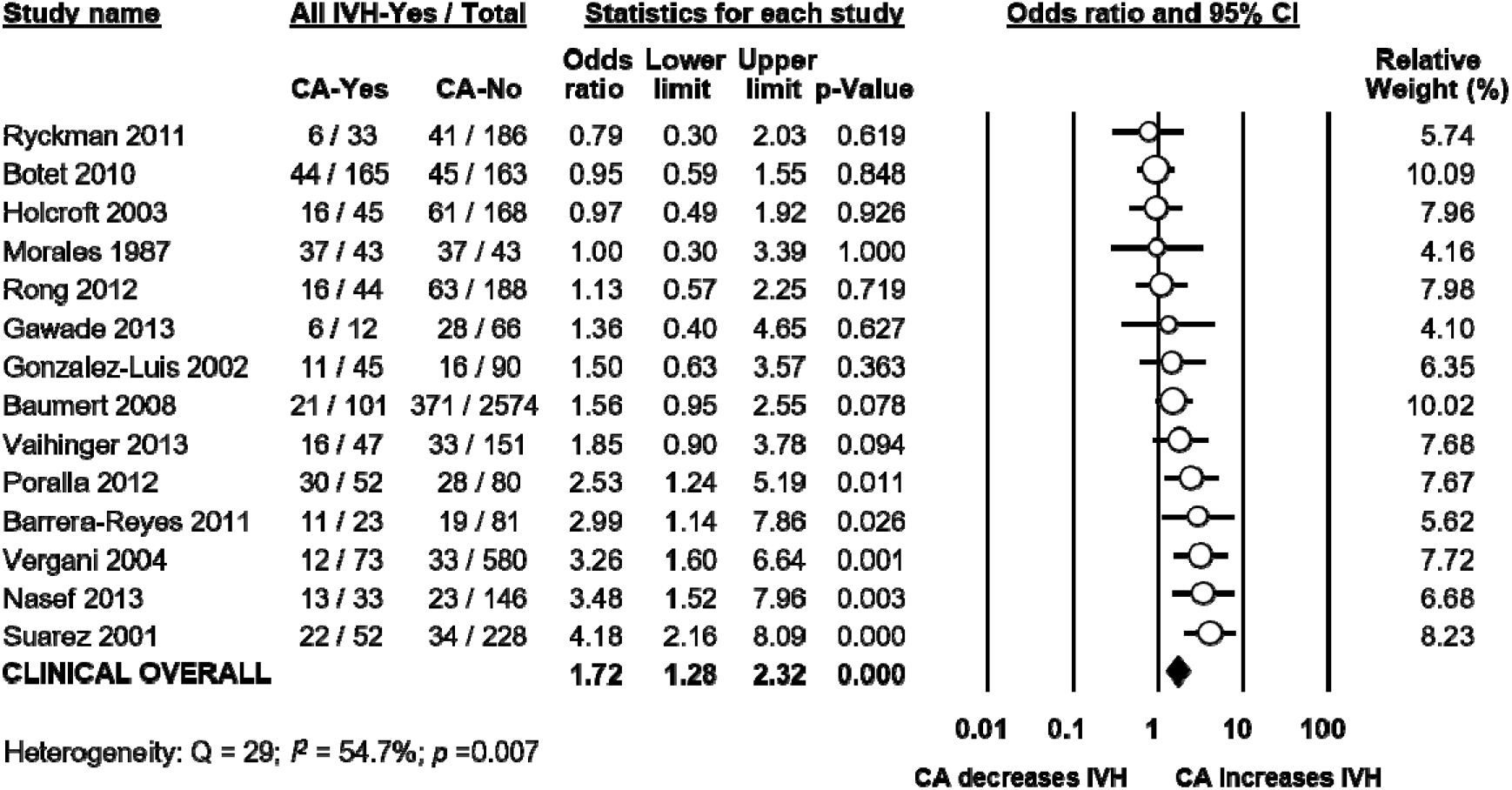
Meta-analysis of the association between clinical chorioamnionitis (CA) and all grades intraventricular hemorrhage (IVH).

**Figure 6.**
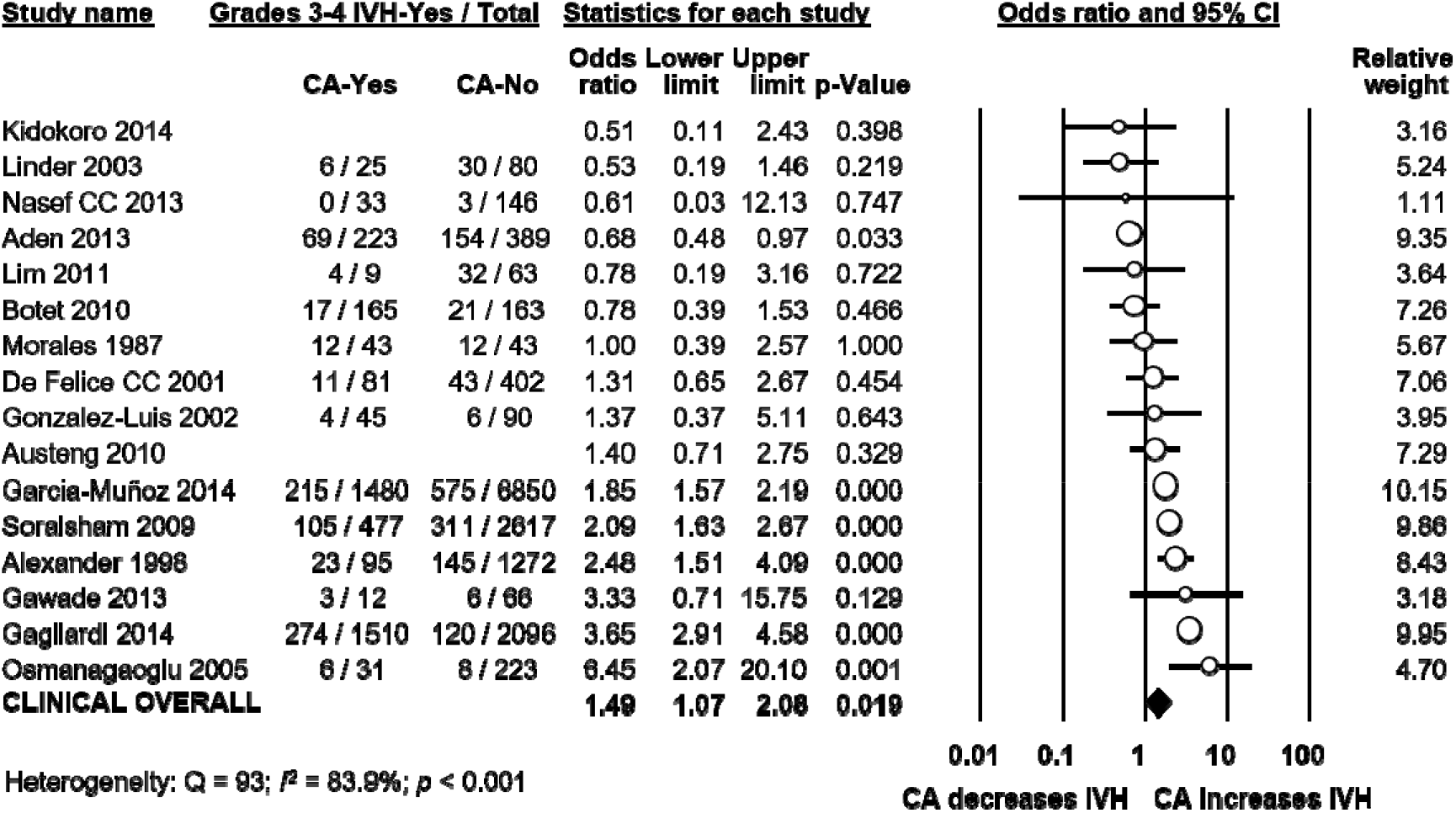
Meta-analysis of the association between clinical chorioamnionitis (CA) and grades 3-4 intraventricular hemorrhage (IVH). CI: confidence interval.

**Figure 7.**
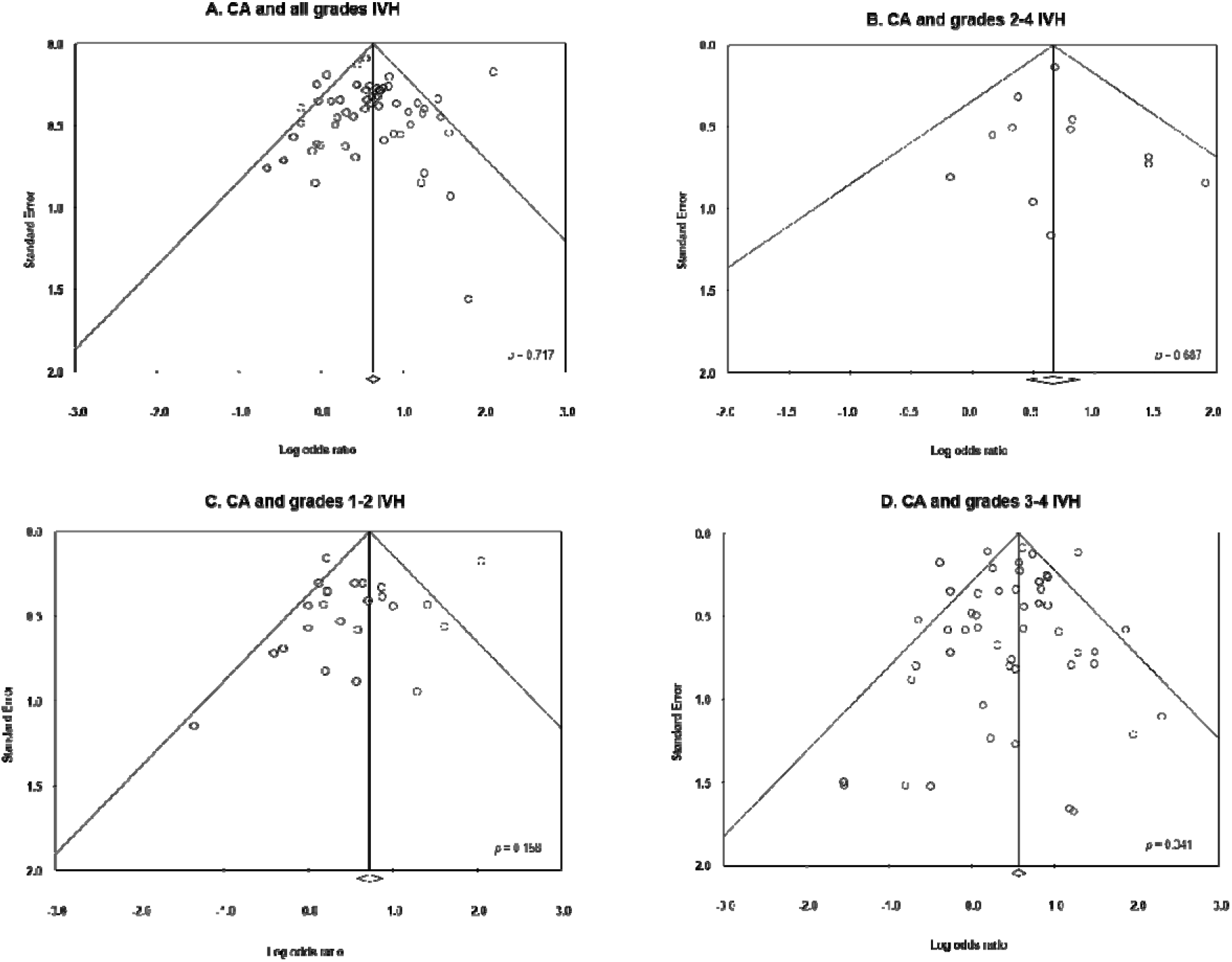
Funnel plots assessing publication bias for the association between chorioamnionitis (CA) and intraventricular hemorrhage (IVH).

### 3.4 Analysis of covariates

To confirm findings from earlier reports (20, 22) on the differences in baseline and clinical characteristics between infants with and without CA, we carried out further meta-analyses. Infants exposed to CA had significantly lower GA and BW, as shown in Table 1. Moreover, infants with CA had significantly higher rates of exposure to ACS, significantly higher rates of PROM, significantly higher rates of EOS, significantly higher rates of LOS, and significantly higher rates of PDA (Table 1). Infants with CA also had significantly lower rates of cesarean delivery, significantly lower rates of small for gestational age (SGA) and significantly lower rates of preeclampsia (Table 1).

**Table 1.** Meta-analysis of the association between chorioamnionitis and covariates.

| Meta-analysis | Chorioamni<br>onitis | *k* | Effect size | 95% CI | *Z* | *p* | Heterogeneity |  |  |
| --- | --- | --- | --- | --- | --- | --- | --- | --- | --- |
|  |  |  |  |  |  |  | *Q* | *p* | *I*<sup>2</sup> |
| Gestational age<br>(weeks) | Clinical | 11 | MD -0.73 | -1.16 to -0.30 | -3.35 | 0.001 | 140 | <0.001 | 92.9 |
|  | Histological | 42 | MD -1.27 | -1.49 to -1.05 | -11.42 | <0.001 | 495 | <0.001 | 91.7 |
|  | Any type | 56 | MD -1.20 | -1.40 to -1.00 | -11.66 | <0.001 | 839 | <0.001 | 93.4 |
| Birth weight (g) | Clinical | 11 | MD -29.14 | -77.66 to 19.39 | -1.18 | 0.239 | 107 | <0.001 | 90.6 |
|  | Histological | 41 | MD -70.21 | -96.71 to -43.72 | -5.19 | <0.001 | 362 | <0.001 | 89.0 |
|  | Any type | 55 | MD -55.00 | -74.89 to -35.12 | -5.42 | <0.001 | 474 | <0.001 | 88.6 |
| Antenatal<br>corticosteroids | Clinical | 5 | OR 1.10 | 0.76 to 1.60 | 0.52 | 0.605 | 55 | <0.001 | 92.8 |
|  | Histological | 31 | OR 1.20 | 1.01 to 1.42 | 2.03 | 0.043 | 95 | <0.001 | 68.3 |
|  | Any type | 38 | OR 1.19 | 1.02 to 1.38 | 2.28 | 0.023 | 155 | <0.001 | 76.1 |
| Cesarean<br>section | Clinical | 8 | OR 0.53 | 0.30 to 0.93 | -2.23 | 0.026 | 316 | <0.001 | 97.8 |
|  | Histological | 28 | OR 0.33 | 0.25 to 0.45 | -7.32 | <0.001 | 177 | <0.001 | 84.8 |
|  | Any type | 36 | OR 0.37 | 0.29 to 0.47 | -8.06 | <0.001 | 495 | <0.001 | 92.9 |
| SGA | Histological | 15 | OR 0.33 | 0.23 to 0.49 | -5.53 | <0.001 | 79 | <0.001 | 82.3 |
|  | Any type | 16 | OR 0.34 | 0.23 to 0.50 | -5.64 | <0.001 | 80 | <0.001 | 81.2 |
| Preeclampsia | Clinical | 3 | OR 0.16 | 0.09 to 0.29 | -5.98 | <0.001 | 2 | 0.369 | <0.001 |
|  | Histological | 23 | OR 0.15 | 0.11 to 0.20 | -13.09 | <0.001 | 65 | <0.001 | 66.2 |
|  | Any type | 27 | OR 0.15 | 0.12 to 0.20 | -15.25 | <0.001 | 69 | <0.001 | 62.2 |
| PROM | Clinical | 3 | OR 5.02 | 2.71 to 9.31 | 5.12 | <0.001 | 2 | <0.001 | 18 |
|  | Histological | 27 | OR 3.14 | 2.54 to 3.87 | 10.63 | <0.001 | 149 | <0.001 | 82.6 |
|  | Any type | 30 | OR 3.29 | 2.70 to 4.02 | 11.76 | <0.001 | 155 | <0.001 | 81.3 |
| Male sex | Clinical | 8 | OR 1.10 | 0.80 to 1.53 | 0.58 | 0.560 | 80 | <0.001 | 91.2 |
|  | Histological | 35 | OR 0.99 | 0.89 to 1.11 | -0.15 | 0.881 | 91 | <0.001 | 62.8 |
|  | Any type | 46 | OR 1.00 | 0.90 to 1.12 | 0.07 | 0.941 | 193 | <0.001 | 76.7 |
| Maternal<br>diabetes | Any type | 9 | OR 0.81 | 0.65 to 1.01 | -1.92 | 0.055 | 5 | 0.725 | 0.0 |
| EOS | Clinical | 7 | OR 4.41 | 3.58 to 5.42 | 14.08 | <0.001 | 9 | 0.197 | 30.3 |
|  | Histological | 18 | OR 2.62 | 1.88 to 3.65 | 5.68 | <0.001 | 48 | <0.001 | 64.9 |
|  | Any type | 25 | OR 3.81 | 3.20 to 4.54 | 14.96 | <0.001 | 87 | <0.001 | 72.4 |
| LOS | Clinical | 5 | OR 1.41 | 1.10 to 1.81 | 2.68 | 0.007 | 17 | 0.002 | 76.8 |
|  | Histological | 30 | OR 1.53 | 1.27 to 1.84 | 4.45 | <0.001 | 134 | <0.001 | 78.3 |
|  | Any type | 37 | OR 1.55 | 1.34 to 1.80 | 5.82 | <0.001 | 174 | <0.001 | 79.3 |
| PDA | Clinical | 4 | OR 1.30 | 1.04 to 1.64 | 2.27 | 0.023 | 7 | 0.062 | 59.0 |
|  | Histological | 26 | OR 1.41 | 1.15 to 1.72 | 3.35 | 0.001 | 144 | <0.001 | 82.6 |
|  | Any type | 31 | OR 1.60 | 1.35 to 1.80 | 6.00 | <0.001 | 195 | <0.001 | 84.6 |
| RDS | Clinical | 3 | OR 2.01 | 0.48 to 8.41 | 0.95 | 0.341 | 11 | 0.004 | 82.0 |
|  | Histological | 15 | OR 1.09 | 0.81 to 1.45 | 0.55 | 0.582 | 89 | <0.001 | 84.3 |
|  | Any type | 21 | OR 1.00 | 0.78 to 1.29 | 0.01 | 0.990 | 149 | <0.001 | 86.6 |

CI: confidence interval; MD: mean difference; OR: odds ratio; SGA: small for gestational age; PROM: premature rupture of membranes; EOS: early-onset sepsis; LOS: late-onset sepsis; PDA: patent ductus arteriosus; RDS: respiratory distress syndrome.

We carried out meta-regression analysis to determine the possible influence of GA and BW on the association between CA and IVH. As Table 2 shows, meta-regression did not find that differences in GA or BW had a significant effect on the association between CA and IVH.

**Table 2.**
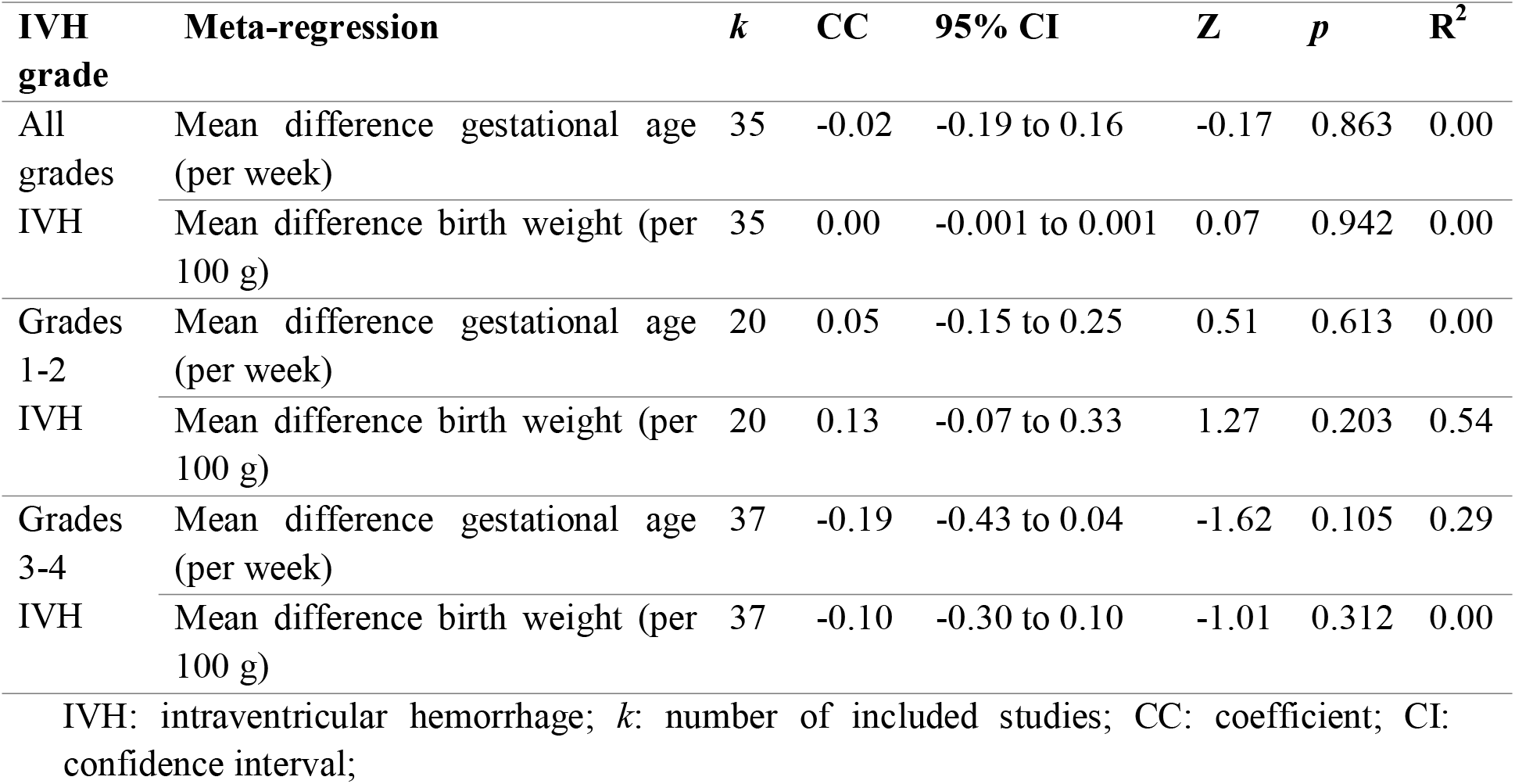
Random effects meta-regression of IVH risk in the chorioamnionitis group, and mean difference in gestational age and birth weight.

To further analyze the effect of GA on the risk of IVH, we carried out subgroup analyses. We found that in the group of studies where the CA-group did not differ significantly (p>0.05) in GA from the control group, CA was a risk factor for all grades IVH but not for grades 3-4 IVH (Table 4). We analyzed a subgroup of studies where the CA-group had a MD in GA of ≤0.5 weeks, and we found that CA was a risk factor for all grades IVH and for grades 3-4 IVH (Table 4). We also found a that in studies where the CA-group had a MD in GA of less than 1 week, CA was a risk factor for all grades IVH and for grades 3-4 IVH (Table 4).

**Table 3.** Random effects meta-regression of IVH risk in the chorioamnionitis group, and predefined covariates.

| IVH grade | Meta-regression | k | CC | 95% CI | Z | P | R<sup>2</sup> |
| --- | --- | --- | --- | --- | --- | --- | --- |
| All grades IVH | Chorioamnionitis type (histological/clinical) | 49 | -0.08 | -0.45 to 0.29 | -0.42 | 0.673 | 0.00 |
|  | Funisitis (CA+F+ vs. CA+F-) | 9 | -0.12 | -0.62 to 0.38 | -0.49 | 0.627 | 0.00 |
|  | ACS (log OR) | 25 | 0.01 | -0.46 to 0.49 | 0.05 | 0.964 | 0.00 |
|  | Cesarean section (log OR) | 22 | -0.07 | -0.47 to 0.32 | -0.36 | 0.717 | 0.00 |
|  | Maternal age (MD) | 17 | -0.08 | -0.24 to 0.07 | -1.09 | 0.276 | 0.00 |
|  | SGA (log OR) | 11 | 0.39 | -0.09 to 0.88 | 1.59 | 0.111 | 0.54 |
|  | PROM (log OR) | 20 | -0.40 | -1.06 to 0.26 | -1.19 | 0.233 | 0.08 |
|  | Preeclampsia (log OR) | 16 | 0.03 | -0.40 to 0.45 | 0.13 | 0.897 | 0.00 |
|  | Mortality (log OR) | 26 | 0.18 | -0.15 to 0.50 | 1.07 | 0.285 | 0.07 |
|  | Early onset sepsis (log OR) | 14 | 0.13 | -0.47 to 0.73 | 0.42 | 0.672 | 0.00 |
|  | Late onset sepsis (log OR) | 24 | -0.04 | -0.39 to 0.32 | -0.19 | 0.846 | 0.00 |
|  | PDA (log OR) | 22 | 0.13 | -0.21 to 0.48 | 0.75 | 0.453 | 0.00 |
|  | RDS (log OR) | 21 | 0.02 | -0.22 to 0.27 | 0.22 | 0.827 | 0.00 |
| Grades 3-4 IVH | Chorioamnionitis type (histological/clinical) | 46 | -0.07 | -0.44 to 0.30 | -0.37 | 0.708 | 0.00 |
|  | Funisitis (CA+F+ vs. CA+F-) | 8 | 0.07 | -0.29 to 0.44 | 0.40 | 0.691 | 0.00 |
|  | ACS (log OR) | 28 | -0.11 | -0.60 to 0.38 | 0.42 | 0.672 | 0.00 |
|  | Cesarean section (log OR) | 27 | 0.08 | -0.09 to 0.26 | 0.92 | 0.358 | 0.06 |
|  | Maternal age (MD) | 18 | -0.01 | -0.25 to 0.22 | -0.11 | 0.911 | 0.00 |
|  | SGA (log OR) | 12 | 0.24 | -0.19 to 0.67 | 1.08 | 0.280 | 0.22 |
|  | PROM (log OR) | 22 | -0.09 | -0.35 to 0.17 | -0.65 | 0.515 | 0.00 |
|  | Preeclampsia (log OR) | 18 | 0.41 | 0.20 to 0.63 | 3.74 | 0.0004 | 1.00 |
|  | Mortality (log OR) | 30 | 0.42 | 0.17 to 0.67 | 3.33 | 0.001 | 0.58 |
|  | Early onset sepsis (log OR) | 20 | 0.11 | -0.32 to 0.54 | 0.51 | 0.613 | 0.00 |
|  | Late onset sepsis (log OR) | 26 | 0.35 | 0.01 to 0.70 | 1.99 | 0.047 |  |
|  | PDA (log OR) | 21 | 0.40 | 0.04 to 0.76 | 2.18 | 0.029 | 0.48 |
|  | RDS (log OR) | 27 | -0.06 | -0.34 to 0.21 | -0.49 | 0.627 | 0.14 |

k: number of studies included; CA: chorioamnionitis; F: funisitis; CC: coefficient; CI: confidence interval; MD: mean difference; OR: odds ratio; ACS: antenatal corticosteroids; SGA: small for gestational age; PROM: premature rupture of membranes; PDA: patent ductus arteriosus; RDS: respiratory distress syndrome.

**Table 4.** Subgroup meta-analyses based on difference in gestational age (GA).

| Subgroup of studies | IVH definition | k | OR | 95% CI | *p* |
| --- | --- | --- | --- | --- | --- |
| Studies where CA-group did not differ significantly in GA from control ($p>0.05$) | All grades IVH | 15 | 1.59 | 1.20 to 2.10 | 0.001 |
|  | Grades 3-4 IVH | 14 | 1.54 | 0.99 to 2.39 | 0.055 |
| Studies where CA-group did differ significantly in GA from control ($p<0.05$) | All grades IVH | 20 | 1.71 | 1.46 to 2.01 | <0.001 |
|  | Grades 3-4 IVH | 23 | 1.90 | 1.56 to 2.33 | 0.000 |
| Studies where CA-group had a MD in GA of $\leq 0.5$ weeks compared to control | All grades IVH | 11 | 1.66 | 1.25 to 2.22 | <0.001 |
|  | Grades 3-4 IVH | 13 | 1.36 | 1.03 to 1.80 | 0.028 |
| Studies where CA-group had a MD in GA of $>0.5$ weeks compared to control | All grades IVH | 24 | 1.68 | 1.43 to 1.98 | <0.001 |
|  | Grades 3-4 IVH | 23 | 1.99 | 1.64 to 2.41 | <0.001 |
| Studies where CA-group had a MD in GA of $<1$ weeks compared to control | All grades IVH | 16 | 1.72 | 1.37 to 2.15 | <0.001 |
|  | Grades 3-4 IVH | 18 | 1.52 | 1.21 to 1.92 | <0.001 |
| Studies where CA-group had a MD in GA of $\geq 1$ weeks compared to control | All grades IVH | 19 | 1.65 | 1.37 to 1.97 | <0.001 |
|  | Grades 3-4 IVH | 18 | 2.00 | 1.60 to 2.50 | <0.001 |

CA: chorioamnionitis; GA: gestational age; IVH: intraventricular hemorrhage; k: number of studies included; CI: confidence interval; MD: mean difference.

To evaluate the role of other prespecified covariates in the association between CA and IVH, we performed additional meta-regression analyses. Meta-regression could not find a significant difference in IVH risk between infants with clinical and infants with histological CA (Table 3). Meta-regression did find a significant association between the CA-associated risk of grades 3-4 IVH and the risk of preeclampsia (Supplementary Figure 1), mortality (Supplementary Figure 2), risk of LOS (Supplementary Figure 3) and risk of PDA (Supplementary Figure 4) Other meta-regressions could not find a significant association between the CA-associated risk of IVH and other covariates (Table 3).

### 3.5 Analysis of funisitis

To evaluate the role of funisitis (i.e. fetal inflammatory response) in the development of IVH, we carried out further meta-analyses. Thirteen studies reported on IVH (35, 38, 60, 64, 69, 70, 73, 81, 87, 88, 97, 102, 103) in infants with histological CA with or without funisitis. As shown in Figure 8, meta-analysis could not show a significant difference in IVH risk between infants with funisitis and infants with CA without funisitis (OR all grades IVH: 1.22, 95% CI 0.89 to 1.67; grades 3-4 IVH: 1.17, 95% CI 0.74 to 1.85). Using meta-regression, we also found no significant difference in IVH risk between infants with funisitis, and infants with CA without funisitis (Table 3).

**Figure 8.**
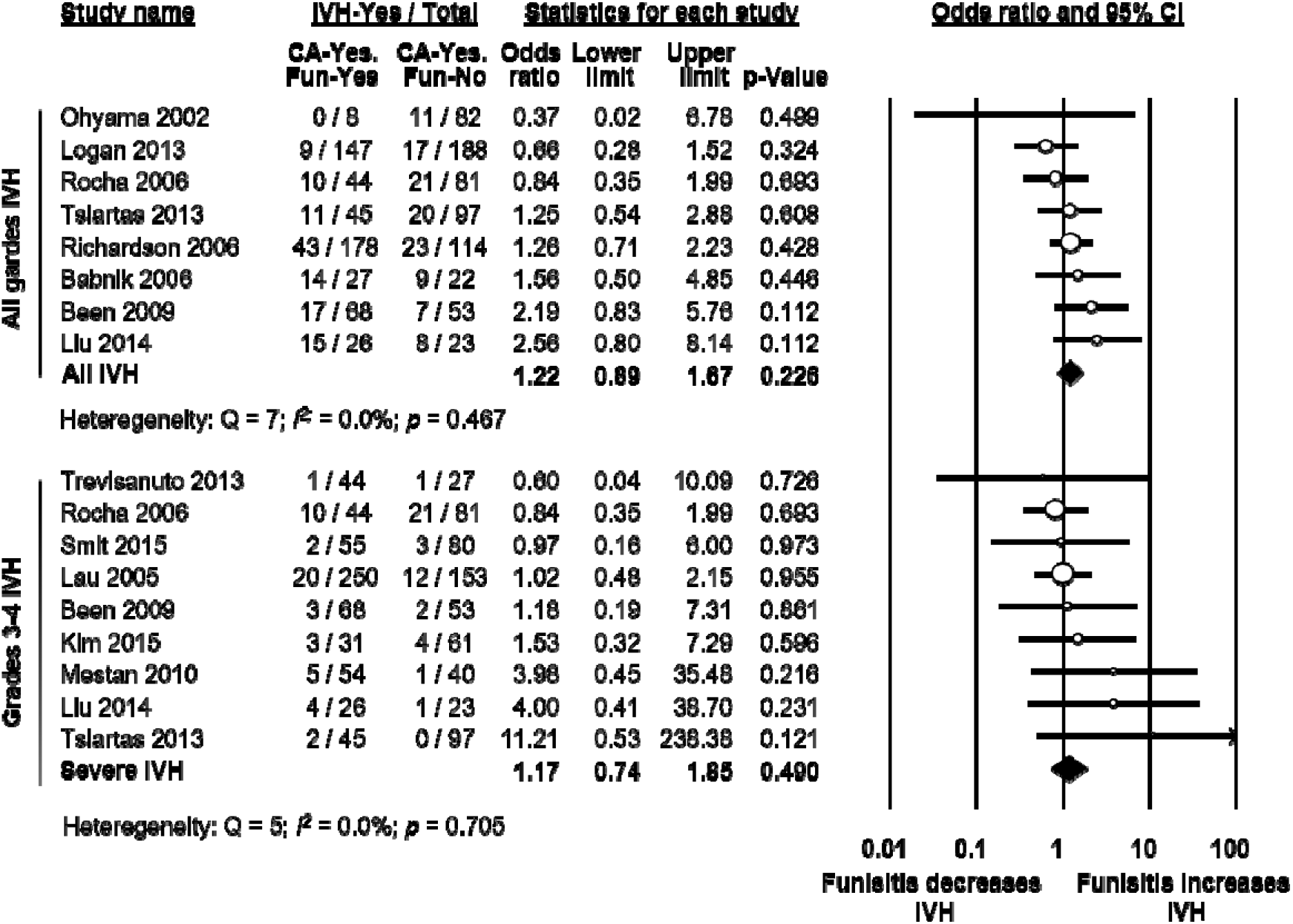
Meta-analysis of the association between funisitis and intraventricular hemorrhage (IVH). Fun: funisitis; CI: confidence interval.

### 3.6 Analysis based on adjusted data

Thirteen studies adjusted the association between CA and the risk of IVH for confounding factors. As shown in Supplementary Table 3 and 4, studies adjusted for different covariates. Meta-analysis pooling this adjusted data found that CA was significantly associated with a higher risk of all grades IVH (OR 1.25, 95% CI 1.02 to 1.53, Supplementary Table 3). This association became non-significant when only analyzing studies which used a histological definition of CA (Supplementary Table 3). Meta-analysis of adjusted data also found a significant association between CA and grades 3-4 IVH (OR 1.22, 95% CI 1.04 to 1.43, Supplementary Table 4). This association became non-significant when grouping studies by clinical or histological CA definition (Supplementary Table 4).

## 4. Discussion

The current systematic review and meta-analysis demonstrates that both clinical and histological CA are associated with an increased risk for developing IVH in very preterm infants. In contrast, the presence of funisitis did not increase IVH risk when compared to CA in the absence of funisitis. We found through additional meta-analyses that CA-exposed infants had significantly lower GA and BW than infants not exposed to CA. However, metaregression and subgroup analysis could not demonstrate an association between the lower GA and BW and the risk of IVH in the CA-exposed infants. This suggests that the effects of CA on HIV risk might be at least partially independent on the role of CA as an etiological factor for very preterm birth.

The association between CA and increased risk of HIV is biologically and clinically plausible. IVH generally occurs within the three first days of life and affects the infants with higher hemodynamic and respiratory instability, frequently associated with extreme prematurity and/or severe perinatal infections (2, 115). Therefore, the clinical circumstances around birth and during the first days of life are critical for the development of IVH. Our study confirms previous reports showing that these clinical circumstances are different in CA-exposed and CA–unexposed very preterm infants (17, 20, 22). Thus, CA-exposed infants were born 1.2 weeks earlier, they were 55g lighter at birth, and they were more frequently exposed to ACS, PROM, vaginal delivery, early and late onset sepsis, and PDA. As mentioned in the introduction, some of these factors may have affected IVH risk.

The degree of prematurity is the most important predisposing factor for the occurrence of IVH (1–3), as well as for other complications of prematurity such as BPD, ROP, NEC, or PDA (20, 22, 23, 116, 117). Nevertheless, very preterm birth is always a pathological event and very preterm infants have a morbidity and mortality risk associated with whichever condition led to their early delivery (118–121). Therefore, CA may affect infant morbidity through inducing very preterm birth or through the deleterious effects of infection/inflammation. Interestingly, previous meta-analyses showed an association between the lower gestational age of the CA-exposed group and the CA-associated risk of BPD (116), PDA (20), and ROP (22, 23). In contrast, our meta-regression could not show that the difference in GA between CA-exposed and CA-unexposed infants significantly correlated with IVH risk. Moreover, we performed subgroup analyses in which we only included the studies showing small or no differences in GA between the CA-exposed and the control group and we observed that the significant IVH risk was maintained in this subgroup of studies. In contrast, this was not the case when the same subgroup analysis was performed for PDA (20) or ROP (22, 23). Altogether this suggests that CA may increase complications such as PDA or ROP through GA-dependent mechanisms, whereas the effect on IVH may be mediated by GA-independent mechanisms.

Besides GA, several other factors potentially confound the association between CA and IVH. A number of studies provided data adjusted for confounding factors, but confounders accounted for in each model differed across studies. We performed separate analyses aggregating adjusted association measures. This reduced or made non-significant the association between CA and IVH (Supplementary Table 3 and 4). Earlier meta-analyses on the association between CA and cerebral palsy (18), BPD (17), ROP (22) also showed that the positive association found when unadjusted data were pooled, was reduced or became nonsignificant when only adjusted data were pooled. Moreover, in our previous meta-analysis on CA and PDA (20) we found that CA was risk factor for PDA when unadjusted data were pooled, and that CA was a protective factor for PDA when adjusted data were pooled.

Adjustment for potential confounders, particularly for GA and/or BW, is a common strategy used in observational studies analyzing predictors of outcomes of prematurity (120). Quality assessment tools such as the NOS even downgrade studies for not adjusting for confounding factors. However, adjustment for GA and BW is controversial and can arguably lead to biased conclusions (118, 120). Preterm infants are at risk of adverse outcomes both due to their immaturity and due to the pathological conditions that led to their preterm birth (118–120, 122). Very low GA is therefore both a risk factor for adverse outcomes, as well as a mediator in the causal pathway that links preterm birth to adverse outcomes (118, 120). The problem with adjusting for intermediate variables, such as GA, is that it may introduce bias unless each confounder is accounted for in the model (118, 120, 123, 124). As discussed by Gagliardi et al., “the difficulty of achieving—at least at the current level of knowledge of etiology of preterm birth—full control of all mediator–outcome confounders limits the possibility of causal interpretation of the associations found but not their descriptive value” (121, p. 798). In this sense, by providing analysis of both the unadjusted and adjusted data, our study may be a valuable contribution to the understanding of CA as etiopathogenic factor of both prematurity and IVH.

Our data suggest that CA-exposed infants are not only younger but also more clinically unstable than the non-exposed infants. This is reflected in the higher mortality and the higher rate of sepsis and PDA in CA-exposed infants (Table 1). Of note, meta-regression showed a correlation between the effect size of the association between CA and grade 3-4 IVH and the effect sizes of the association between CA and PDA. As mentioned elsewhere, the presence of a hemodynamically relevant PDA has been correlated with the occurrence of IVH and the proposed mechanism is the disturbance of cerebral blood flow (2, 3, 8, 125). Our data support this association between IVH and PDA in CA-exposed infants.

The biological plausibility of the association between CA and IVH is supported by the direct and indirect effects of inflammatory mediators. Hemodynamic disturbances in preterm infants with CA have been correlated with elevated cord blood concentrations of proinflammatory cytokines such as IL-6, IL-1beta and TNF-alfa (126). Cytokines can act directly on the vascular smooth muscle, producing vascular relaxation and hypotension or indirectly by increasing the production of endothelium-derived vasoactive mediators (126). In addition, cytokines can eventually promote a neuro-inflammatory cascade in the fetal brain penetrating across the blood brain barrier or activating specific receptors such as CD14 and TLR4 which are constitutively expressed in the circumventricular organs, choroid plexus and leptomeninges (127, 128). Inflamed glial or endothelial cells, challenged by external stimuli, enhance the release/expression of various chemoattractants and adhesion molecules which may promote the platelet and neutrophil activation and adhesion determining possible endothelial cell damage and changes in blood rheology and flow (129, 130). These changes, occurring inside the fragile germinal matrix capillaries or within the vascular connection between germinal matrix and the subependymal venous network, may increase the likelihood of IVH in preterm infants with CA.

We have discussed the role of funisitis in earlier meta-analyses on CA and ROP (22) and CA and PDA (20). It is worth noting that not all intraamniotic infections will induce an inflammatory response in the fetus (131). Funisitis is generally considered the histologic counterpart to fetal inflammatory response syndrome (16, 132). We found that exposure to funisitis did not significantly increase the risk of IVH, when compared to exposure to CA without funisitis. This is an argument against role of the fetal inflammatory response in the etiopathogenesis of IVH. We have previously reported that funisitis is not an additional risk factor for developing PDA (20) but the presence of funisitis significantly increased the risk of developing severe ROP (22).

Our meta-analysis has several limitations that should be considered. Firstly, there was substantial heterogeneity in how CA was defined in studies. The definitions of clinical CA in particular varied substantially, and recent recommendations propose restricting the term CA to pathologic diagnosis (133). Secondly, only 5 out of 85 included studies studied the association between CA and IVH as their main objective. However, this could also have reduced the effect of publication bias. Thirdly, only 13 out of the 85 included studies provided adjusted data, and they used different models and adjusted for different confounders. Finally, in this study and earlier studies on ROP (22) and PDA (20), we had a much more limited number of studies to draw from for analyzing funisitis than when analyzing CA. The strengths of our study include: the use of a comprehensive search, duplication of screening, inclusion and data extraction to reduce bias, a large number of included studies, and an extensive analysis of confounding factors, through meta-analysis, meta-regression and the inclusion of adjusted data.

A significant limitation in any meta-analysis on IVH is the potential for heterogeneity in defining the condition. The grading system most commonly used for neonatal IVH was first reported by Papile et al. and later modified by Volpe and is based on the presence and amount of blood in the germinal matrix, lateral ventricles, and periventricular white matter (1). Grade 1 represents germinal matrix hemorrhage only with no or minimal IVH (<10% of ventricular area on parasagittal view). When IVH occupies 10–50% of ventricular area on parasagittal view, it is defined as grade 2 (1). Grade 3 is IVH with blood occupying more than 50% of the ventricular area on parasagittal view. Grade 4 represents severe IVH with associated periventricular echodensity (1). Although grade 4 IVH is a periventricular hemorrhagic infarction rather than an extension of IVH per se, the 1-4 grading system remains pervasive in the literature and clinical setting despite debate regarding appropriate nomenclature (134). In addition grade 3 and 4 IVHs are frequently grouped together as severe or high grade IVH (134). Nevertheless, our meta-analysis shows a significant increased risk of both severe and less severe (grade 1-2) IVH in CA-exposed infants. Therefore, potential differences in IVH classification may not have affected the results.

## 5. Conclusion

IVH is a multifactorial complication that is more common in more preterm and more clinically unstable infants. We established for the first time through meta-analysis that CA is a risk factor for IVH. We also confirmed earlier findings that CA is a risk factor for being born more preterm and presenting more clinical instability. However, in contrast to other complications of prematurity, such as PDA, ROP, or BPD (20, 22, 23, 116), the effect of CA on IVH appears to be independent of CA as a causative factor for very preterm birth.

## 6. Conflict of Interest

The authors declare that the research was conducted in the absence of any commercial or financial relationships that could be construed as a potential conflict of interest.

## 7. Author Contributions

E.V.-M. carried out data collection, carried out statistical analyses, assessed methodological quality, contributed to interpretation of results, drafted the initial manuscript, and reviewed and revised the manuscript. M.F. contributed to the design of the study, the statistical analysis and interpretation of results and reviewed and revised the manuscript. O. M. R. selected studies for inclusion, carried out data collection and carried out statistical analyses. S.P. contributed to interpretation of results and reviewed and revised the manuscript. G.C. contributed to interpretation of results and reviewed and revised the manuscript. P.D. carried out and supervised data collection and contributed to interpretation of results. F.M. contributed to interpretation of results and reviewed and revised the manuscript. E.V. conceptualized and designed the study, carried out the search and selected studies for inclusion, supervised data collection, contributed to statistical analyses and interpretation of results, and reviewed and revised the manuscript. All authors approved the final manuscript as submitted.

## 8. Data Availability Statement

The datasets generated and analyzed for this study can be found in the Harvard Dataverse (135): https://dataverse.harvard.edu/dataset.xhtml?persistentId=doi%3A10.7910%2FDVN%2FJ9RHUF

## References

1. Volpe JJ. Impaired neurodevelopmental outcome after mild germinal matrix-intraventricular hemorrhage. Pediatrics. 2015:peds. 2015–3553.

2. Inder TE, Perlman JM, Volpe JJ. Preterm Intraventricular Hemorrhage/Posthemorrhagic Hydrocephalus. Volpe’s Neurology of the Newborn (Sixth Edition): Elsevier; 2018. p. 637–98.e21.

3. Ballabh P. Intraventricular hemorrhage in premature infants: mechanism of disease. Pediatric research. 2010;67(1):1.

4. Tortora D, Severino M, Malova M, Parodi A, Morana G, Sedlacik J, et al. Differences in subependymal vein anatomy may predispose preterm infants to GMH–IVH. Archives of Disease in Childhood-Fetal and Neonatal Edition. 2017:fetalneonatal-2017-312710.

5. Romantsik O, Bruschettini M, Calevo MG, Banzi R, Ley D. Pharmacological pain and sedation interventions for the prevention of intraventricular hemorrhage in preterm infants on assisted ventilation □an overview of systematic reviews. The Cochrane Library. 2017.

6. Ramenghi LA, Fumagalli M, Groppo M, Consonni D, Gatti L, Bertazzi PA, et al. Germinal matrix hemorrhage: intraventricular hemorrhage in very-low-birth-weight infants: the independent role of inherited thrombophilia. Stroke. 2011;42(7):1889–93.

7. Bermick J, Dechert R, Sarkar S. Does hyperglycemia in hypernatremic preterm infants increase the risk of intraventricular hemorrhage? Journal of Perinatology. 2016;36(9):729.

8. Poryo M, Boeckh JC, Gortner L, Zemlin M, Duppré P, Ebrahimi-Fakhari D, et al. Ante-, peri-and postnatal factors associated with intraventricular hemorrhage in very premature infants. Early human development. 2018;116:1–8.

9. Chevallier M, Debillon T, Pierrat V, Delorme P, Kayem G, Durox M, et al. Leading causes of preterm delivery as risk factors for intraventricular hemorrhage in very preterm infants: results of the EPIPAGE 2 cohort study. American Journal of Obstetrics & Gynecology. 2017;216(5):518.e1–.e12.

10. DiSalvo D. The correlation between placental pathology and intraventricular hemorrhage in the preterm infant. The Developmental Epidemiology Network Investigators. Pediatric research. 1998;43(1):15–9.

11. Jackson CM, Wells CB, Tabangin ME, Meinzen-Derr J, Jobe AH, Chougnet CA. Proinflammatory immune responses in leukocytes of premature infants exposed to maternal chorioamnionitis or funisitis. Pediatric research. 2017;81(2):384.

12. Cornette L. Fetal and neonatal inflammatory response and adverse outcome. Semin Fetal Neonatal Med. 2004;9(6):459–70.

13. Thomas W, Speer CP. Chorioamnionitis: important risk factor or innocent bystander for neonatal outcome? Neonatology. 2011;99(3):177–87.

14. Tita AT, Andrews WW. Diagnosis and management of clinical chorioamnionitis. Clin Perinatol. 2010;37(2):339–54.

15. Pugni L, Pietrasanta C, Acaia B, Merlo D, Ronchi A, Ossola MW, et al. Chorioamnionitis and neonatal outcome in preterm infants: a clinical overview. J Matern Fetal Neonatal Med. 2015:1–5.

16. Gantert M, Been JV, Gavilanes AW, Garnier Y, Zimmermann LJ, Kramer BW. Chorioamnionitis: a multiorgan disease of the fetus? J Perinatol. 2010;30 Suppl:S21–30.

17. Hartling L, Liang Y, Lacaze-Masmonteil T. Chorioamnionitis as a risk factor for bronchopulmonary dysplasia: a systematic review and meta-analysis. Arch Dis Child Fetal Neonatal Ed. 2012;97(1):F8–F17.

18. Wu YW, Colford JM, Jr. Chorioamnionitis as a risk factor for cerebral palsy: A metaanalysis. Jama. 2000;284(11):1417–24.

19. Park HW, Choi YS, Kim KS, Kim SN. Chorioamnionitis and Patent Ductus Arteriosus: A Systematic Review and Meta-Analysis. PLoS One. 2015;10(9):e0138114.

20. Behbodi E, Villamor-Martínez E, Degraeuwe PL, Villamor E. Chorioamnionitis appears not to be a Risk Factor for Patent Ductus Arteriosus in Preterm Infants: A Systematic Review and Meta-Analysis. Scientific reports. 2016;6.

21. Been JV, Lievense S, Zimmermann LJ, Kramer BW, Wolfs TG. Chorioamnionitis as a risk factor for necrotizing enterocolitis: a systematic review and meta-analysis. J Pediatr. 2013;162(2):236–42e2.

22. Villamor-Martinez E, Cavallaro G, Raffaeli G, Rahim OMM, Gulden S, Ghazi AM, et al. Chorioamnionitis as a risk factor for retinopathy of prematurity: an updated systematic review and meta-analysis. bioRxiv. 2018:291–476.

23. Mitra S, Aune D, Speer CP, Saugstad OD. Chorioamnionitis as a risk factor for retinopathy of prematurity: a systematic review and meta-analysis. Neonatology. 2014;105(3):189–99.

24. Strunk T, Inder T, Wang X, Burgner D, Mallard C, Levy O. Infection-induced inflammation and cerebral injury in preterm infants. The Lancet infectious diseases. 2014;14(8):751–62.

25. Moher D, Liberati A, Tetzlaff J, Altman DG, Group P. Preferred reporting items for systematic reviews and meta-analyses: the PRISMA statement. PLoS med. 2009;6(7):e1000097.

26. Wan X, Wang W, Liu J, Tong T. Estimating the sample mean and standard deviation from the sample size, median, range and/or interquartile range. BMC medical research methodology. 2014;14(1):135.

27. Borenstein M, Hedges LV, Higgins J, Rothstein HR. Subgroup analyses. Introduction to Meta-analysis2009. p. 149–86.

28. Borenstein M, Hedges LV, Higgins J, Rothstein HR. Identifying and quantifying heterogeneity. Introduction to Meta-Analysis2009. p. 107–26.

29. Borenstein M, Hedges LV, Higgins J, Rothstein HR. Meta □Regression. Introduction to Meta-Analysis2009. p. 187–203.

30. Ådén U, Lin A, Carlo W, Leviton A, Murray JC, Hallman M, et al. Candidate gene analysis: severe intraventricular hemorrhage in inborn preterm neonates. Journal of pediatrics. 2013;163(5):1503–6.e1.

31. Ahn HM, Park EA, Cho SJ, Kim Y-J, Park H-S. The association of histological chorioamnionitis and antenatal steroids on neonatal outcome in preterm infants born at less than thirty-four weeks’ gestation. Neonatology. 2012;102(4):259–64.

32. Alexander JM, Gilstrap LC, Cox SM, McIntire DM, Leveno KJ. Clinical chorioamnionitis and the prognosis for very low birth weight infants. Obstetrics & Gynecology. 1998;91(5, Part 1):725–9.

33. Arayici S, Kadioglu Simsek G, Oncel MY, Eras Z, Canpolat FE, Oguz SS, et al. The effect of histological chorioamnionitis on the short-term outcome of preterm infants≤ 32 weeks: a single-center study. Journal of Maternal-Fetal & Neonatal Medicine. 2014;27(11):1129–33.

34. Austeng D, Blennow M, Ewald U, Fellman V, Fritz T, Hellstrom-Westas L, et al. Incidence of and risk factors for neonatal morbidity after active perinatal care: extremely preterm infants study in Sweden (EXPRESS). Acta Paediatrica. 2010;99(7):978–92.

35. Babnik J, Stucin-Gantar I, Kornhauser-Cerar L, Sinkovec J, Wraber B, Derganc M. Intrauterine inflammation and the onset of peri-intraventricular hemorrhage in premature infants. Neonatology. 2006;90(2):113–21.

36. Barrera-Reyes R, Ruiz-Macias H, Segura-Cervantes E. [Neurodevelopment at one year of age [corrected] in preterm newborns with history of maternal chorioamnionitis]. Ginecologia y obstetricia de Mexico. 2011;79(1):31–7.

37. Baumert M, Brozek G, Paprotny M, Walencka Z, Sodowska H, Cnota W, et al. Epidemiology of peri/intraventricular haemorrhage in newborns at term. J Physiol Pharmacol. 2008;59(Suppl 4):67–75.

38. Been JV, Rours IG, Kornelisse RF, Passos VL, Kramer BW, Schneider TA, et al. Histologic chorioamnionitis, fetal involvement, and antenatal steroids: effects on neonatal outcome in preterm infants. American journal of obstetrics and gynecology. 2009;201(6):587.e1–.e8.

39. Bordigato MA, Piva D, Di Gangi IM, Giordano G, Chiandetti L, Filippone M. Asymmetric dimethylarginine in ELBW newborns exposed to chorioamnionitis. Early human development. 2011;87(2):143–5.

40. Botet F, Figueras J, Carbonell-Estrany X, Arca G, Group CS. Effect of maternal clinical chorioamnionitis on neonatal morbidity in very-low birthweight infants: a case-control study. Journal of perinatal medicine. 2010;38(3):269–73.

41. Bry K, Jacobsson B, Nilsson S, Bry K. Gastric fluid cytokines are associated with chorioamnionitis and white blood cell counts in preterm infants. Acta Paediatrica. 2015;104(6):575–80.

42. Dalton J, Dechert RE, Sarkar S. Assessment of association between rapid fluctuations in serum sodium and intraventricular hemorrhage in hypernatremic preterm infants. American journal of perinatology. 2015;32(08):795–802.

43. Dempsey E, Chen M-F, Kokottis T, Vallerand D, Usher R. Outcome of neonates less than 30 weeks gestation with histologic chorioamnionitis. American journal of perinatology. 2005;22(03):155–9.

44. Dexter SC, Pinar H, Malee MP, Hogan J, Carpenter MW, Vohr BR. Outcome of very low birth weight infants with histopathologic chorioamnionitis. Obstetrics & Gynecology. 2000;96(2):172–7.

45. Ecevit A, Anuk-İnce D, Yapakçı E, Kupana-Ayva Ş, Kurt A, Yanık FF, et al. Association of respiratory distress syndrome and perinatal hypoxia with histologic chorioamnionitis in preterm infants. Turkish Journal of Pediatrics. 2014;56(1).

46. Elimian A, Verma U, Beneck D, Cipriano R, Visintainer P, Tejani N. Histologic chorioamnionitis, antenatal steroids, and perinatal outcomes. Obstetrics & Gynecology. 2000;96(3):333–6.

47. Erdemir G, Kultursay N, Calkavur S, Zekioğlu O, Koroglu OA, Cakmak B, et al. Histological chorioamnionitis: effects on premature delivery and neonatal prognosis. Pediatrics & Neonatology. 2013;54(4):267–74.

48. Fung G, Bawden K, Chow P, Yu V. Long-term Outcome of Extremely Preterm Infants Following Chorioamnionitis 絨毛膜羊膜炎對極早早產兒的長遠影響. HK J Paediatr (new series). 2003;8(2):87–92.

49. Gagliardi L, Rusconi F, Bellù R, Zanini R, Network IN. Association of maternal hypertension and chorioamnionitis with preterm outcomes. Pediatrics. 2014;134(1):e154–e61.

50. Garcia-Munoz Rodrigo F, Galan Henriquez G, Figueras Aloy J, Garcia-Alix Perez A, Network S. Outcomes of very-low-birth-weight infants exposed to maternal clinical chorioamnionitis: a multicentre study. Neonatology. 2014;106(3):229–34.

51. Gawade PL, Whitcomb BW, Chasan-Taber L, Pekow PS, Ronnenberg AG, Shah B, et al. Second stage of labor and intraventricular hemorrhage in early preterm infants in the vertex presentation. Journal of Maternal-Fetal & Neonatal Medicine. 2013;26(13): 1292–8.

52. González-Luis G, García IJ, Rodríguez-Miguélez J, Mussons FB, Aloy JF, editors. Patología neonatal en los menores de 1.500 gramos con relación al antecedente de corioamnionitis. Anales de Pediatría; 2002. Included study Elsevier.

53. Gray PH, Hurley TM, Rogers YM, O’Callaghan MJ, Tudehope DI, Burns YR, et al. Survival and neonatal and neurodevelopmental outcome of 24-29 week gestation infants according to primary cause of preterm delivery. Australian and New Zealand journal of obstetrics and gynaecology. 1997;37(2):161–8.

54. Hendson L, Russell L, Robertson CM, Liang Y, Chen Y, Abdalla A, et al. Neonatal and neurodevelopmental outcomes of very low birth weight infants with histologic chorioamnionitis. Journal of Pediatrics. 2011;158(3):397–402.

55. Hitti J, Tarczy-Hornoch P, Murphy J, Hillier SL, Aura J, Eschenbach DA. Amniotic fluid infection, cytokines, and adverse outcome among infants at 34 weeks’ gestation or less. Obstetrics & Gynecology. 2001;98(6):1080–8.

56. Holcroft CJ, Blakemore KJ, Allen M, Graham EM. Association of prematurity and neonatal infection with neurologic morbidity in very low birth weight infants. Obstetrics & Gynecology. 2003;101(6):1249–53.

57. Kallankari H, Kaukola T, Ojaniemi M, Herva R, Perhomaa M, Vuolteenaho R, et al. Chemokine CCL18 predicts intraventricular hemorrhage in very preterm infants. Annals of medicine. 2010;42(6):416–25.

58. Kaukola T, Herva R, Perhomaa M, Pääkkö E, Kingsmore S, Vainionpää L, et al. Population cohort associating chorioamnionitis, cord inflammatory cytokines and neurologic outcome in very preterm, extremely low birth weight infants. Pediatric research. 2006;59(3):478–83.

59. Kidokoro H, Anderson PJ, Doyle LW, Woodward LJ, Neil JJ, Inder TE. Brain injury and altered brain growth in preterm infants: predictors and prognosis. Pediatrics. 2014:peds. 2013–336.

60. Kim SY, Choi CW, Jung E, Lee J, Lee JA, Kim H, et al. Neonatal Morbidities Associated with Histologic Chorioamnionitis Defined Based on the Site and Extent of Inflammation in Very Low Birth Weight Infants. Journal of Korean medical science. 2015;30(10):1476–82.

61. Kirchner L, Helmer H, Heinze G, Wald M, Brunbauer M, Weninger M, et al. Amnionitis with Ureaplasma urealyticum or other microbes leads to increased morbidity and prolonged hospitalization in very low birth weight infants. Eur J Obstet Gynecol Reprod Biol. 2007;134(1):44–50.

62. Klebermass-Schrehof K, Czaba C, Olischar M, Fuiko R, Waldhoer T, Rona Z, et al. Impact of low-grade intraventricular hemorrhage on long-term neurodevelopmental outcome in preterm infants. Child’s nervous system. 2012;28(12):2085–92.

63. Kosuge S, Ohkuchi A, Minakami H, Matsubara S, Uchida A, Eguchi Y, et al. Influence of chorioamnionitis on survival and morbidity in singletons live-born at< 32 weeks of gestation. Acta obstetricia et gynecologica Scandinavica. 2000;79(10):861–5.

64. Lau J, Magee F, Qiu Z, Houbé J, Von Dadelszen P, Lee SK. Chorioamnionitis with a fetal inflammatory response is associated with higher neonatal mortality, morbidity, and resource use than chorioamnionitis displaying a maternal inflammatory response only. American journal of obstetrics and gynecology. 2005;193(3):708–13.

65. Lee HJ, Kim E-K, Kim H-S, Choi CW, Kim BI, Choi J-H. Chorioamnionitis, respiratory distress syndrome and bronchopulmonary dysplasia in extremely low birth weight infants. Journal of Perinatology. 2011;31(3):166–70.

66. Lee JY, Kim HS, Jung E, Kim ES, Shim GH, Lee HJ, et al. Risk factors for periventricular-intraventricular hemorrhage in premature infants. Journal of Korean medical science. 2010;25(3):418–24.

67. Lim W, Lien R, Chiang M, Fu R, Lin J, Chu S, et al. Hypernatremia and grade III/IV intraventricular hemorrhage among extremely low birth weight infants. Journal of Perinatology. 2011; 31(3):193–8.

68. Linder N, Haskin O, Levit O, Klinger G, Prince T, Naor N, et al. Risk factors for intraventricular hemorrhage in very low birth weight premature infants: a retrospective case-control study. Pediatrics. 2003;111(5):e590–e5.

69. Liu Z, Tang Z, Li J, Yang Y. Effects of placental inflammation on neonatal outcome in preterm infants. Pediatrics & Neonatology. 2014;55(1):35–40.

70. Logan JW, Westra SJ, Allred EN, O’Shea TM, Kuban K, Paneth N, et al. Antecedents of perinatal cerebral white matter damage with and without intraventricular hemorrhage in very preterm newborns. Pediatric neurology. 2013;49(2):88–96.

71. Lu H, Wang Q, Lu J, Zhang Q, Kumar P. Risk factors for intraventricular hemorrhage in preterm infants born at 34 weeks of gestation or less following preterm premature rupture of membranes. Journal of Stroke and Cerebrovascular Diseases. 2016;25(4):807–12.

72. Mehta R, Nanjundaswamy S, Shen-Schwarz S, Petrova A. Neonatal morbidity and placental pathology. Indian journal of pediatrics. 2006;73(1):25–8.

73. Mestan K, Yu Y, Matoba N, Cerda S, Demmin B, Pearson C, et al. Placental inflammatory response is associated with poor neonatal growth: preterm birth cohort study. Pediatrics. 2010;125(4):e891–e8.

74. Miyazaki K, Furuhashi M, Ishikawa K, Tamakoshi K, Hayashi K, Kai A, et al. Impact of chorioamnionitis on short-and long-term outcomes in very low birth weight preterm infants: the Neonatal Research Network Japan. Journal of Maternal-Fetal & Neonatal Medicine. 2016;29(2):331–7.

75. Morales WJ. The effect of chorioamnionitis on the developmental outcome of preterm infants at one year. Obstetrics & Gynecology. 1987;70(2):183–6.

76. Mu S-C, Lin C-H, Chen Y-L, Ma H-J, Lee J-S, Lin M-I, et al. Impact on neonatal outcome and anthropometric growth in very low birth weight infants with histological chorioamnionitis. Journal of the Formosan Medical Association. 2008;107(4):304–10.

77. Nasef N, Shabaan AE, Schurr P, Iaboni D, Choudhury J, Church P, et al. Effect of clinical and histological chorioamnionitis on the outcome of preterm infants. American journal of perinatology. 2013;30(01):059–68.

78. Ogunyemi D, Murillo M, Jackson U, Hunter N, Alperson B. The relationship between placental histopathology findings and perinatal outcome in preterm infants. Journal of Maternal-Fetal & Neonatal Medicine. 2003;13(2):102–9.

79. Oh KJ, Park JY, Lee J, Hong J-S, Romero R, Yoon BH. The combined exposure to intra-amniotic inflammation and neonatal respiratory distress syndrome increases the risk of intraventricular hemorrhage in preterm neonates. Journal of perinatal medicine. 2018;46(1):9–20.

80. Oh S-H, Kim J-j, Do H-j, Lee BS, Kim K-S, Kim EA-R. Preliminary Study on Neurodevelopmental Outcome and Placental Pathology among Extremely Low Birth Weight Infants. Korean J Perinatol. 2015;26(1):67–77.

81. Ohyama M, Itani Y, Yamanaka M, Goto A, Kato K, Ijiri R, et al. Re-evaluation of chorioamnionitis and funisitis with a special reference to subacute chorioamnionitis. Human pathology. 2002;33(2):183–90.

82. Osmanağaoğlu MA, Ünal S, Bozkaya H. Chorioamnionitis risk and neonatal outcome in preterm premature rupture of membranes. Archives of gynecology and obstetrics. 2005;271(1):33–9.

83. Pappas A, Kendrick DE, Shankaran S, Stoll BJ, Bell EF, Laptook AR, et al. Chorioamnionitis and early childhood outcomes among extremely low-gestational-age neonates. JAMA Pediatr. 2014;168(2):137–47.

84. Perrone S, Toti P, Toti MS, Badii S, Becucci E, Gatti MG, et al. Perinatal outcome and placental histological characteristics: a single-center study. Journal of Maternal-Fetal & Neonatal Medicine. 2012;25(sup1): 110–3.

85. Polam S, Koons A, Anwar M, Shen-Schwarz S, Hegyi T. Effect of chorioamnionitis on neurodevelopmental outcome in preterm infants. Archives of pediatrics & adolescent medicine. 2005;159(11):1032–5.

86. Poralla C, Hertfelder H-J, Oldenburg J, Müller A, Bartmann P, Heep A. Elevated interleukin-6 concentration and alterations of the coagulation system are associated with the development of intraventricular hemorrhage in extremely preterm infants. Neonatology. 2012;102(4):270–5.

87. Richardson BS, Wakim E, Walton J. Preterm histologic chorioamnionitis: impact on cord gas and pH values and neonatal outcome. American journal of obstetrics and gynecology. 2006;195(5):1357–65.

88. Rocha G, Proença E, Quintas C, Rodrigues T, Guimarães H. Chorioamnionitis and neonatal morbidity. Acta Medica Portuguesa. 2006;19(3):207–12.

89. Rodríguez J Trujillo A, Cobo T, Vives I, Bosch J, Kacerovsky M, Posadas DE, et al. Gestational age is more important for short term neonatal outcome than microbial invasion of the amniotic cavity or intra Jamniotic inflammation in preterm prelabor rupture of membranes. Acta obstetricia et gynecologica Scandinavica. 2016;95(8):926–33.

90. Rong Z, Liu H, Xia S, Chang L. Risk and protective factors of intraventricular hemorrhage in preterm babies in Wuhan, China. Child’s Nervous System. 2012;28(12):2077–84.

91. Ryckman KK, Dagle JM, Kelsey K, Momany AM, Murray JC. Replication of genetic associations in the inflammation, complement, and coagulation pathways with intraventricular hemorrhage in LBW preterm neonates. Pediatric research. 2011;70(1):90–5.

92. Salas AA, Faye-Petersen OM, Sims B, Peralta-Carcelen M, Reilly SD, McGwin G, et al. Histological characteristics of the fetal inflammatory response associated with neurodevelopmental impairment and death in extremely preterm infants. Journal of Pediatrics. 2013;163(3):652–7.e2.

93. Sarkar S, Kaplan C, Wiswell TE, Spitzer AR. Histological chorioamnionitis and the risk of early intraventricular hemorrhage in infants born ≤ 28 weeks gestation. Journal of perinatology. 2005;25(12):749–52.

94. Sato M, Nishimaki S, Yokota S, Seki K, Horiguchi H, An H, et al. Severity of chorioamnionitis and neonatal outcome. Journal of Obstetrics and Gynaecology Research. 2011;37(10):1313–9.

95. Seliga-Siwecka JP, Kornacka MK. Neonatal outcome of preterm infants born to mothers with abnormal genital tract colonisation and chorioamnionitis: a cohort study. Early human development. 2013;89(5):271–5.

96. Shankaran S, Lin A, Maller-Kesselman J, Zhang H, O’shea TM, Bada HS, et al. Maternal race, demography, and health care disparities impact risk for intraventricular hemorrhage in preterm neonates. Journal of Pediatrics. 2014;164(5):1005–11.e3.

97. Smit AL, Been JV, Zimmermann LJ, Kornelisse RF, Andriessen P, Vanterpool SF, et al. Automated auditory brainstem response in preterm newborns with histological chorioamnionitis. Journal of Maternal-Fetal & Neonatal Medicine. 2015;28(15):1864–9.

98. Soraisham A, Trevenen C, Wood S, Singhal N, Sauve R. Histological chorioamnionitis and neurodevelopmental outcome in preterm infants. Journal of Perinatology. 2013;33(1):70–5.

99. Soraisham AS, Singhal N, McMillan DD, Sauve RS, Lee SK, Network CN. A multicenter study on the clinical outcome of chorioamnionitis in preterm infants. American journal of obstetrics and gynecology. 2009;200(4):372.e1–.e6.

100. Suarez RD, Grobman WA, Parilla BV. Indomethacin tocolysis and intraventricular hemorrhage. Obstetrics & Gynecology. 2001;97(6):921–5.

101. Suppiej A, Franzoi M, Vedovato S, Marucco A, Chiarelli S, Zanardo V. Neurodevelopmental outcome in preterm histological chorioamnionitis. Early human development. 2009;85(3):187–9.

102. Trevisanuto D, Peruzzetto C, Cavallin F, Vedovato S, Cosmi E, Visentin S, et al. Fetal placental inflammation is associated with poor neonatal growth of preterm infants: a case-control study. Journal of Maternal-Fetal & Neonatal Medicine. 2013;26(15):1484–90.

103. Tsiartas P, Kacerovsky M, Musilova I, Hornychova H, Cobo T, Sävman K, et al. The association between histological chorioamnionitis, funisitis and neonatal outcome in women with preterm prelabor rupture of membranes. Journal of Maternal-Fetal & Neonatal Medicine. 2013;26(13):1332–6.

104. Vaihinger M, Mazzitelli N, Balanian N, Grandi C. The relationship between placental lesions and early hemorrhagic-ischemic cerebral injury in very low birth weight infants. Rev Fac Cienc Med Cordoba. 2012;70(3):123–33.

105. van Vliet EO, de Kieviet JF, van der Voorn JP, Been JV, Oosterlaan J, van Elburg RM. Placental pathology and long-term neurodevelopment of very preterm infants. American journal of obstetrics and gynecology. 2012;206(6):489.e1–.e7.

106. Vergani P, Locatelli A, Doria V, Assi F, Paterlini G, Pezzullo JC, et al. Intraventricular hemorrhage and periventricular leukomalacia in preterm infants. Obstetrics & Gynecology. 2004;104(2):225–31.

107. Watterberg KL, Gerdes JS, Gifford KL, Lin H-M. Prophylaxis against early adrenal insufficiency to prevent chronic lung disease in premature infants. Pediatrics. 1999;104(6):1258–63.

108. Wirbelauer J, Thomas W, Speer CP. Response of leukocytes and nucleated red blood cells in very low-birth weight preterm infants after exposure to intrauterine inflammation. Journal of Maternal-Fetal & Neonatal Medicine. 2011;24(2):348–53.

109. Xu L, Ren R, Zhu S, Zhuang H, Huang Z, Yang H. Effect of chorioamnionitis on brain injury in preterm infants. Zhongguo dang dai er ke za zhi. 2012;14(9):661–3.

110. Yamada N, Sato Y, Moriguchi-Goto S, Yamashita A, Kodama Y, Sameshima H, et al. Histological severity of fetal inflammation is useful in predicting neonatal outcome. Placenta. 2015;36(12): 1490–3.

111. Yanowitz TD, Potter DM, Bowen AD, Baker RW, Roberts JM. Variability in cerebral oxygen delivery is reduced in premature neonates exposed to chorioamnionitis. Pediatric research. 2006;59(2):299–304.

112. Yoon BH, Romero R, Kim CJ, Jun JK, Gomez R, Choi J-H, et al. Amniotic fluid interleukin-6: a sensitive test for antenatal diagnosis of acute inflammatory lesions of preterm placenta and prediction of perinatal morbidity. American journal of obstetrics and gynecology. 1995;172(3):960–70.

113. Zanardo V, Vedovato S, Suppiej A, Trevisanuto D, Migliore M, Di Venosa B, et al. Histological inflammatory responses in the placenta and early neonatal brain injury. Pediatric and Developmental Pathology. 2008;11(5):350–4.

114. De Felice C, Toti P, Laurini RN, Stumpo M, Picciolini E, Todros T, et al. Early neonatal brain injury in histologic chorioamnionitis. Journal of pediatrics. 2001;138(1):101–4.

115. Mohamed MA, Aly H. Transport of premature infants is associated with increased risk for intraventricular haemorrhage. Archives of Disease in Childhood-Fetal and Neonatal Edition. 2010:fetalneonatal183236.

116. Hartling L, Liang Y, Lacaze-Masmonteil T. Chorioamnionitis as a risk factor for bronchopulmonary dysplasia: a systematic review and meta-analysis. Archives of Disease in Childhood-Fetal and Neonatal Edition. 2012;97(1):F8–F17.

117. Been JV, Lievense S, Zimmermann LJ, Kramer BW, Wolfs TG. Chorioamnionitis as a risk factor for necrotizing enterocolitis: a systematic review and meta-analysis. The Journal of pediatrics. 2013;162(2):236–42.e2.

118. Wilcox AJ, Weinberg CR, Basso O. On the pitfalls of adjusting for gestational age at birth. American journal of epidemiology. 2011;174(9):1062–8.

119. McElrath TF, Hecht JL, Dammann O, Boggess K, Onderdonk A, Markenson G, et al. Pregnancy disorders that lead to delivery before the 28th week of gestation: an epidemiologic approach to classification. American journal of epidemiology. 2008;168(9):980–9.

120. Gagliardi L, Rusconi F, Da Frè M, Mello G, Carnielli V, Di Lallo D, et al. Pregnancy disorders leading to very preterm birth influence neonatal outcomes: results of the population-based ACTION cohort study. Pediatric research. 2013;73(6):794–801.

121. Barros FC, Papageorghiou AT, Victora CG, Noble JA, Pang R, Iams J, et al. The distribution of clinical phenotypes of preterm birth syndrome: implications for prevention. JAMA pediatrics. 2015;169(3):220–9.

122. Basso O, Wilcox A. Mortality risk among preterm babies: Immaturity vs. underlying pathology. Epidemiology (Cambridge, Mass). 2010;21(4):521–7.

123. Basso O, Wilcox A. Mortality risk among preterm babies: immaturity vs. underlying pathology. Epidemiology (Cambridge, Mass). 2010;21(4):521.

124. Hernández-Díaz S, Schisterman EF, Hernán MA. The birth weight “paradox” uncovered? American journal of epidemiology. 2006;164(11):1115–20.

125. Ballabh P. Pathogenesis and prevention of intraventricular hemorrhage. Clinics in perinatology. 2014;41(1):47–67.

126. Yanowitz TD, Jordan JA, Gilmour CH, Towbin R, Bowen AD, Roberts JM, et al. Hemodynamic disturbances in premature infants born after chorioamnionitis: association with cord blood cytokine concentrations. Pediatric research. 2002;51(3):310.

127. McAdams RM, Juul SE. The role of cytokines and inflammatory cells in perinatal brain injury. Neurology research international. 2012;2012.

128. Rivest S. Molecular insights on the cerebral innate immune system. Brain, behavior, and immunity. 2003;17(1):13–9.

129. Molina □Holgado E, Molina □Holgado F. Mending the broken brain: neuroimmune interactions in neurogenesis. Journal of neurochemistry. 2010;114(5):1277–90.

130. Stanimirovic D, Satoh K. Inflammatory mediators of cerebral endothelium: a role in ischemic brain inflammation. Brain Pathology. 2000;10(1):113–26.

131. Revello R, Alcaide MJ, Dudzik D, Abehsera D, Bartha JL. Differential amniotic fluid cytokine profile in women with chorioamnionitis with and without funisitis. The Journal of Maternal-Fetal & Neonatal Medicine. 2015:1–5.

132. Revello R, Alcaide MJ, Dudzik D, Abehsera D, Bartha JL. Differential amniotic fluid cytokine profile in women with chorioamnionitis with and without funisitis. J Matern Fetal Neonatal Med. 2015:1–5.

133. Higgins RD, Saade G, Polin RA, Grobman WA, Buhimschi IA, Watterberg K, et al. Evaluation and management of women and newborns with a maternal diagnosis of chorioamnionitis: summary of a workshop. Obstetrics and gynecology. 2016;127(3):426–36.

134. Leviton A, Kuban K, Paneth N. Intraventricular haemorrhage grading scheme: time to abandon? Acta Pædiatrica. 2007;96(9):1254–6.

135. Chorioamnionitis and Intraventricular Hemorrhage, studies included in systematic review [Internet]. Harvard Dataverse. 2018. Available from: https://doi.org/10.7910/DVN/J9RHUF.

